# The Clonal Hematopoiesis-associated Gene *Srcap* Plays an Essential Role in Hematopoiesis

**DOI:** 10.1101/2024.08.16.607812

**Authors:** Terrence N. Wong, Anna Mychalowych, Ellie R. Feldpausch, Alexander Carson, Darja Karpova, Daniel C. Link

## Abstract

Somatic mutations arising in hematopoietic stem cells (HSCs) may provide the latter with a fitness advantage, allowing the mutant HSC to clonally expand. Such mutations have been recurrently identified in the chromatin modifier, *SRCAP*, in both non-malignant and leukemic clones, suggesting that this gene plays a significant role in hematopoiesis. We generated a conditional *Srcap* loss of function murine model and determined the consequences of hematopoietic-specific loss of this gene. We show that *Srcap* is essential for normal fetal liver erythropoiesis and monocytopoiesis. In *Srcap* deficient fetal livers, the number of phenotypic HSCs is similar to that of controls, but these HSCs exhibit a profound repopulating defect. Likewise, conditional deletion of *Srcap* during adult hematopoiesis results in a rapid loss of HSCs. Loss of *Srcap* is associated with evidence of increased DNA damage in HSCs and lineage-restricted progenitors as assessed by y-H2AX expression. Consistent with this finding, we observed strong transcriptional upregulation of the p53 pathway in *Srcap* deficient erythroid precursors. Collectively our data highlight the importance of *Srcap* in maintaining HSC function and supporting hematopoietic differentiation and suggests that it plays an essential role in maintaining genomic integrity.

**Key Points:**

(1) *Srcap* plays an essential role in supporting normal hematopoietic differentiation. and in maintaining HSC function.
(2) Loss of *Srcap* is associated with evidence of increased DNA damage and transcriptional upregulation of the p53 pathway.

## Introduction

Hematopoietic stem cells (HSCs) acquire somatic mutations with age.^1^ A subset of these mutations, which arise in specific genes, may provide HSCs with a fitness advantage, allowing them to clonally expand in a process termed clonal hematopoiesis (CH).^2,3^ Such mutations can significantly alter the behavior of both HSCs and the maturing hematopoietic populations that are derived from them. These impacts may involve their competitive fitness,^4^ response to cellular stress,^5–7^ leukemogenic potential,^8,9^ inflammatory status,^10^ and interactions with non-hematopoietic tissues^11^ with potential implications in the development of both leukemia^8,9^ and non-hematologic disorders.^12^ As such, understanding the role that genes that are recurrently mutated in CH and malignant leukemias have in hematopoietic populations is of significant biological and clinical importance.

*SRCAP* is a member of the Snf2 family of ATPases.^13^ Along with its ortholog *EP400, SRCAP* is responsible for the incorporation of the variant histone H2A.Z into the nucleosome. H2A.Z incorporation has been associated with multiple intracellular processes including heterochromatin regulation, the DNA damage response, and transcriptional regulation including both activation and repression.^13^ On a cellular level, H2A.Z is essential for the proper functioning of both embryonic^14^ and tissue-specific^15,16^ stem cells, including both their self-renewal and differentiation. Not surprisingly, the dysregulation of H2A.Z has been linked to the development of multiple malignancies.^17,18^ However, the mechanisms underlying how this dysregulation promotes carcinogenesis remain poorly understood. The Srcap complex has also been shown to play an integral role in non-malignant hematopoiesis with hematopoietic-specific loss of *Znhit1*, a component of this complex, impacting both HSC function (quiescence and repopulating ability) and hematopoietic differentiation.^15,19^

We previously assessed for CH in 81 patients who had received prior treatment with cytotoxic therapy.^7^ In addition to mutations in well-characterized CH genes (e.g. *DNMT3A*, *TET2*, *TP53*, *PPM1D*), we identified mutations in *SRCAP* with a variant allele frequency (VAF) > 0.1% in ∼16% of these patients. *SRCAP* mutations have also been observed to arise in CH emerging after allogeneic transplantation^20^ and after treatment for myelodysplastic syndrome (MDS).^21^ In terms of malignant hematologic disease, mutations in *SRCAP* have been identified in 0.9% of MDS cases^22^ and in 5% of core binding factor AML (CBF-AML) cases as a member of the founding clone.^23^ Germline mutations in *SRCAP* have previously been associated with the genetic disorder Floating-Harbor syndrome (FHS), which is characterized by abnormal facial features, skeletal malformations, and delayed speech development.^24^ However, unlike FHS-associated mutations, which typically occur in exons 33-34 of *SRCAP*, CH and AML-associated mutations (which include truncating frameshift and nonsense mutations) have been identified throughout the entire coding region of *SRCAP*. Interestingly, the majority of CBF-AML associated *SRCAP* mutations are truncating in nature.^23^ These findings suggest that *SRCAP* plays a significant role in the regulation of hematopoiesis and that a disruption to SRCAP function may perturb non-malignant hematopoiesis with potential leukemogenic consequences. For these reasons, we assessed how loss of *Srcap* impacts hematopoiesis in mice.

## Materials and methods

### Mouse models

Mice were maintained under standard pathogen-free conditions with animal studies conducted in accordance with IACUC-approved University of Michigan protocols. Using CRISPR/Cas9, the Transgenic, Knockout and Micro-Injection Core at Washington University School of Medicine modified the germline of mice on a C57Bl/6 background, inserting a loxP site before exon 11 and after exon 16 of the *Srcap* gene. Proper insertion of the loxP sites was verified with targeted next-generation sequencing. Vav-iCre (strain 008610) and Mx1-Cre (strain 003556) mice were purchased from The Jackson Laboratory.

For experiments involving adult mice, experimental and control mice were age (8–12 weeks) and sex-matched. For timed mating experiments, 1-3 females were housed with a male for 2 days, separated, and monitored with serial weight checks to assess for pregnancy. Floxing of *Srcap* with the Mx1-Cre strain was induced by the administration of 1.0 µg/g of poly(I:C) (Cytiva) every other day for five total doses as previously described.^25^

### Antibodies

Flow cytometry was performed with the following antibodies (Biolegend unless otherwise noted): Ly-6C/G (RB6-8C5, Gr-1), Ly6G (1A8), Ly6C (HK1.4), CD115 (AFS98), F4/80 (BM8), CD11b (M1/70), Ly-6A/E (D7, Sca1), CD3e (145-2C11), CD16/32 (93, Fc 2R/ Fc 3R), CD34 (RAM34), CD45R (RA3-6B2, B220), CD45.1 (A20, Ly5.1), CD45.2 (104, Ly5.2), CD48 (HM48-1), CD71 (BD Biosciences, C2F2, TfR1), CD117 (2B8, ckit), CD150 (TC15-12F12.2, SLAM), CD201 (ebioscience, 1560, Epcr), CD86 (GL-1), TER-119 (Ter-119), CD44 (IM7), Ki-67(Invitrogen, SolA15), and γ-H2AX (Ser139) (EMD Millipore, JBW301). Cell fixation and permeabilization were performed using the FoxP3 Transcription Factor Fixation/Permeabilization Kit (ebioscience) as per manufacturer’s instructions.

### Flow cytometry

Cells were stained using standard protocols. Flow cytometry data were collected on a 14-color, 3-laser LSRFortessa (BD Biosciences) or a 10-color, 4-laser Gallios (Beckman Coulter) flow cytometer. Cell sorting was performed with a Sony MA900 cell sorter.

### RNA expression

RNA was isolated from cell populations using Trizol LS (Thermo Fisher) as per manufacturer’s instructions. RNA-Seq was performed on ribo-depleted RNA with the library preparation and sequencing performed by the University of Michigan Advanced Genomics Core and analysis performed by the University of Michigan Bioinformatics Core.

### Bone marrow transplantation

For transplantation experiments, Ly5.1/5.2 recipient mice were conditioned with 1,000– 1,100 cGy in split dosing from either a ^137^cesium or x-ray source prior to the retro-orbital transplantation of bone marrow or fetal liver cells. Prophylactic antibiotics (trimethoprim-sulfamethoxazole (Alpharma) or neomycin/polymyxin B (Sigma)) were given during the initial two weeks following transplantation.

### Quantification and statistical analysis

Analysis and quantification of flow cytometry data were performed using FlowJo software (Tree Star). Gene Set Enrichment Analysis (GSEA) of RNA-Seq data was performed with GSEA software,^26^ using the Hallmark gene set annotation pathways. Statistical analysis was done using GraphPad Prism 9 with the specific analytic tests performed detailed in the appropriate figure legends. Unless otherwise noted, data is represented as the mean ± the standard error of the mean.

## Results

### Generation of a mouse model with conditional loss of *Srcap*

In mice, germline homozygous loss of *Srcap* results in early embryonic lethality.^27^ Therefore, to investigate the impact of *Srcap* loss on hematopoiesis, we used CRISPR/Cas9 to generate a conditional *Srcap* loss of function murine model with loxP sites flanking exons 11-16 of the gene (**Figure 1A**). Excision of these exons results in the loss of Srcap’s catalytic ATPase domain and a frameshift mutation. The breeding of *Vav1-iCre^+/−^Srcap^fl/+^*and *Srcap*^fl/fl^ mice together produced no pups with the *Vav1-iCre^+/−^Srcap^fl/fl^* genotype. However, 50 of the first 211 embryos (26 litters) assessed from E13.5 to E16.5 possessed this genotype, consistent with Mendelian genetics. PCR genotyping of sorted fetal liver erythroid progenitor cells from *Vav1-iCre^+/−^Srcap^fl/fl^* embryos showed efficient floxing (**Figure 1B**), while RNA-Seq data from the corresponding fetal liver stage 1 erythroid progenitor cells showed that exons 11-16 were efficiently excised from the *Srcap* mRNA transcript (**Figure 1C**). Thus, *Srcap* is efficiently deleted in hematopoietic cells in *Vav1-iCre^+/−^Srcap^fl/fl^* mice with loss of this gene resulting in embryonic lethality. Consistent with this, *Srcap* deficient E13.5/E14.5 embryos displayed a significantly paler phenotype than their corresponding control littermates with either full expression or haploinsufficiency of *Srcap* (**Figure 1D**).

**Figure 1.**
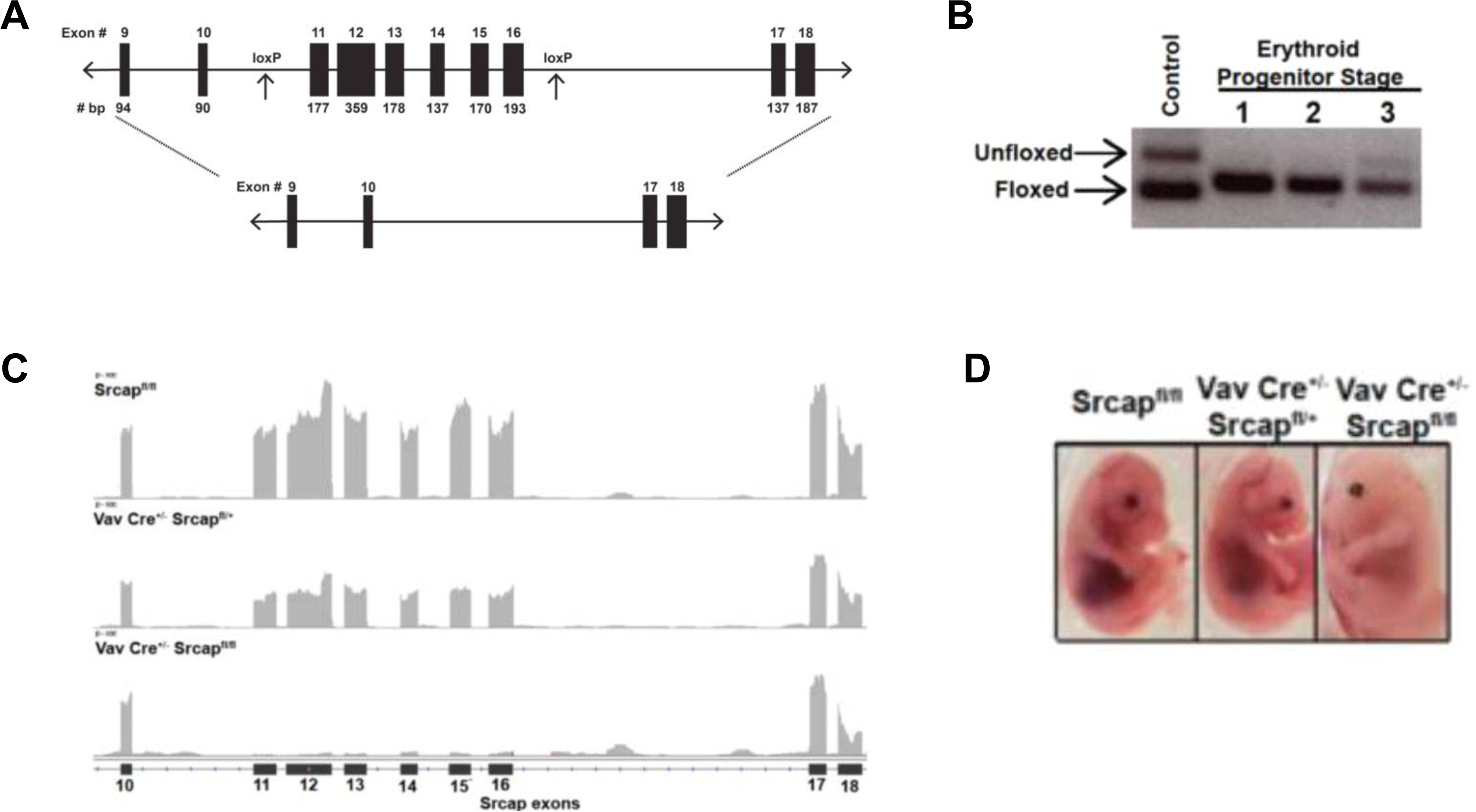
Generation and initial characterization of a mouse model with conditional loss of *Srcap*. (A) Genetic model for conditional loss of *Srcap* in cells expressing Cre recombinase. (B) Representative PCR genotyping reaction showing efficient Cre-induced excision in sorted fetal liver erythroid progenitor cells. (C) Representative RNA-Seq reads of excised *Srcap* exons as displayed by IGV Viewer from sorted fetal liver stage 1 erythroid progenitor cells. (D) Representative pictures of murine embryos with the indicated genotypes.

### Loss of *Srcap* causes aberrant fetal hematopoiesis

We first analyzed how *Srcap* loss impacts mature hematopoietic populations in the fetal liver. Mutations in *SRCAP* have been recurrently identified in myelodysplastic syndrome, suggesting that this gene may play a role in hematopoietic maturation. Consistent with their appearance after H&E staining, *Vav1-iCre^+/−^Srcap^fl/fl^*fetal livers possessed a significantly decreased number of fetal liver cells, with a marked reduction in erythroid Ter119^+^ cells (**Figure 2A-B**, **Supplemental Figure 1A-B**). Interestingly, the number of stage 1 erythroid progenitors (as assessed with CD44 and Ter119 staining along with cell size) was similar between *Vav1-iCre^+/−^ Srcap^fl/fl^* embryos and other genotypes. In contrast, the number of later stage erythroid progenitors was significantly reduced (**Figure 2C-D**, **Supplemental Figure 1C**). Thus, fetal liver erythroid progenitor populations may display differing tolerances to *Srcap* loss, with more mature erythroid populations being particularly sensitive to loss of this gene.

**Figure 2.**
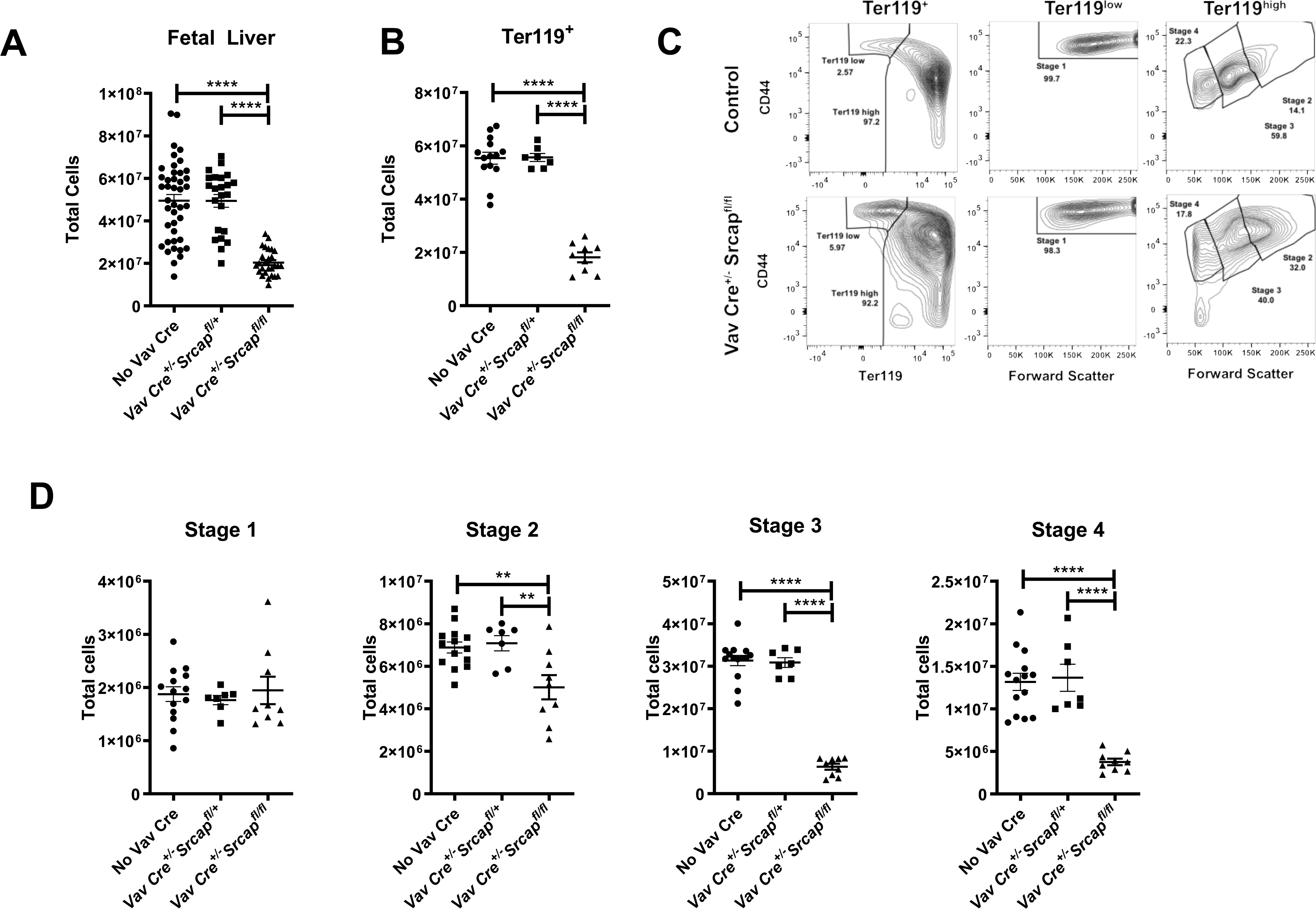
Loss of *Srcap* causes aberrant fetal erythroid maturation. (A) Total number of fetal liver cells. (B) Total number of Ter119^+^ fetal liver cells. (C) Representative flow plots of fetal erythroid maturation. (D) Total number of fetal liver erythroid progenitor cells grouped by their stage of maturation based on forward scatter, Ter119 staining, and CD44 staining. ** P ≤ 0.01. **** P ≤ 0.0001. Significance was determined with an ordinary one-way ANOVA with multiple comparisons.

We also assessed how loss of *Srcap* impacts fetal liver myeloid cell maturation. Although the number of CD11b^+^CD115^−^ phenotypic fetal liver granulocytes was similar in embryos with normal expression, haploinsufficiency, and complete loss of *Srcap,* embryos with complete loss of this gene had significantly fewer phenotypic CD115^+^ monocytes (**Figures 3A-3B; Supplemental Figure 2A**). To ensure that this phenotype was not due to downregulation of CD115, we also used multiple CD115-independent strategies to differentiate between granulocytes and monocytes These included assessing the surface expression of Ly6C (**Figures 3A-3B; Supplemental Figure 2B**), Ly6G, and F4/80 (**Supplemental Figures 2C-E**). Results from these different approaches all demonstrated a marked decline in phenotypic monocytes in concordance with the results obtained with CD115, suggesting that monocytic cells are particularly sensitive to the loss of *Srcap*.

**Figure 3.**
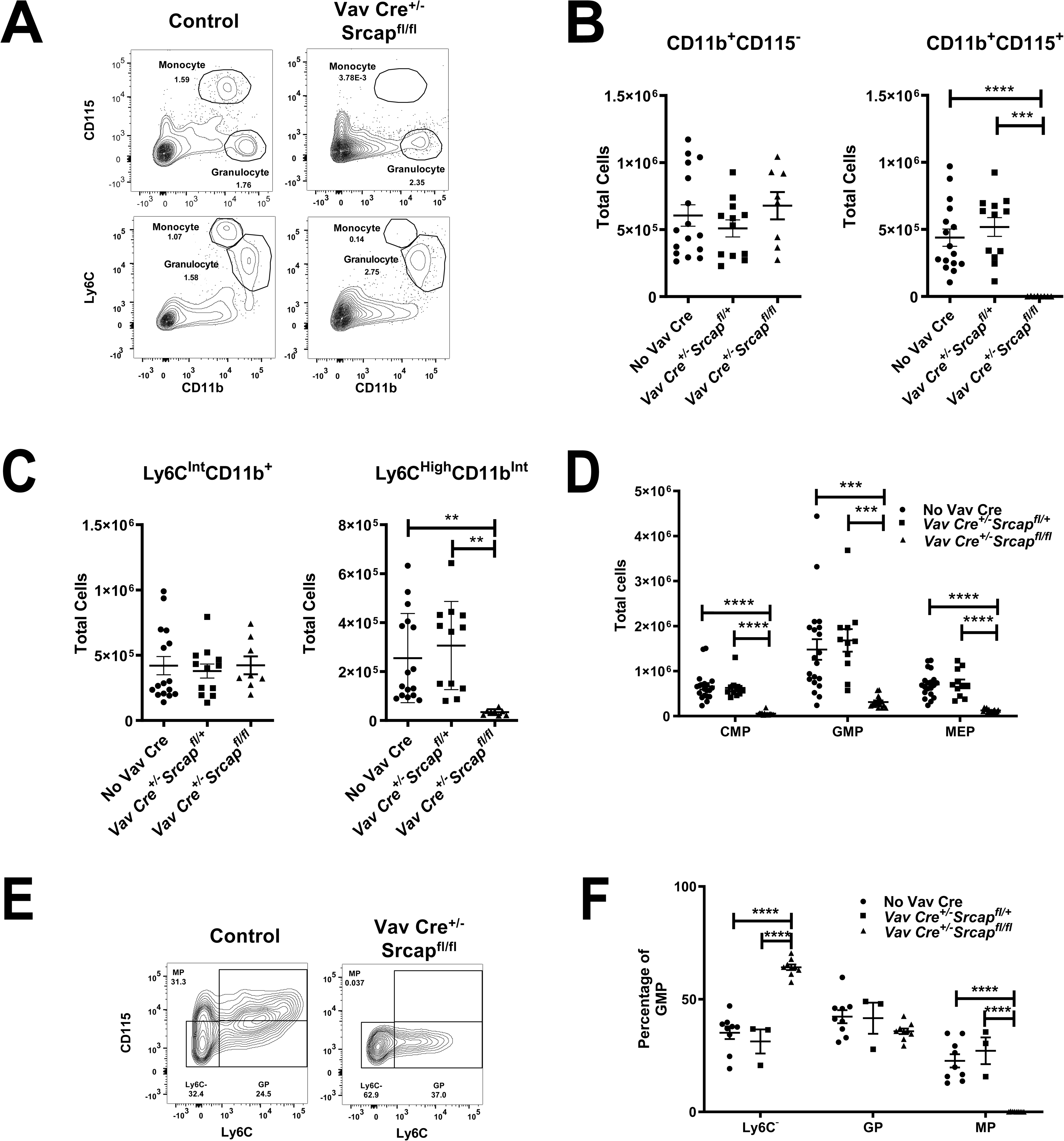
Loss of *Srcap* causes impaired monocytic differentiation. (A) Representative flow plots identifying fetal liver myeloid populations using the markers CD115, Ly6C, and CD11b. (B) Total number of fetal liver cells classified as phenotypic CD11b^+^CD115^−^ granulocytes or CD11b^+^CD115^+^ monocytes as indicated in **A**. (C) Total number of fetal liver cells classified as Ly6C^Int^CD11b^+^ granulocytes or Ly6C^High^CD11b^Int^ monocytes as indicated in **A**. (D) Total number of phenotypic fetal liver common myeloid progenitors (CMPs), granulocyte-monocyte progenitors (GMPs), and megakaryocyte–erythrocyte progenitors (MEPs). (E) Representative flow plots identifying monocytic and granulocytic progenitors within the GMP compartment using Ly6C and CD115 as markers. (F) Percentage of Ly6C^−^ cells, monocyte-committed progenitors (MP), and granulocyte-committed progenitors (GP) within the GMP compartment. ** P ≤ 0.01. *** P ≤ 0.001. **** P ≤ 0.0001. Significance was determined with an ordinary one-way ANOVA with multiple comparisons.

To determine if *Srcap* loss also impacts earlier stages of fetal liver myeloid development, we assessed the number of myeloid progenitors using CD34 and CD16/32 staining. Embryos with loss of *Srcap* had decreased numbers of common myeloid progenitors (CMPs), granulocyte-monocyte progenitors (GMPs), and megakaryocyte-erythrocyte progenitors (MEPs) (**Figure 3D**; **Supplemental Figure 2F**) with a particularly marked decline in CMPs. GMPs have previously been demonstrated to be heterogeneous populations and, using cell surface markers, divisible into granulocyte-biased and monocyte-biased cells.^28^ Consistent with the decrease in mature monocytic populations, the GMP populations of embryos with *Srcap* loss were characterized by a marked loss of monocyte-committed progenitors (MPs) in contrast to more immature Ly6C^−^ cells or granulocytic-committed progenitors (GPs) (**Figure 3E-F**). In total, these data suggest that monocytic populations are particularly sensitive to *Srcap* loss with almost no monocytic-biased progenitors or mature monocytic populations present upon loss of this gene. Collectively, these data show that *Srcap* is required for normal monocytic and erythroid development during fetal hematopoiesis in mice.

### Loss of *Srcap* abrogates fetal HSC function

In addition to MDS, mutations in *SRCAP* have also been recurrently identified in AML. Mutations associated with the development of myeloid malignancies have often been found to provide HSCs with a fitness advantage (e.g. mutations in *DNMT3A* and *TET2*).^4,29^ However, several classes of myeloid malignancy-associated mutations (e.g. impacting the spliceosome or cohesin complexes) have paradoxically been observed to induce defects in HSC function.^30,31^ We therefore analyzed how *Srcap* loss impacts immature hematopoietic populations. In contrast to more differentiated populations, phenotypic CD150^+^CD48^−^ KLS HSCs were present at similar numbers in *Vav1-iCre^+/−^Srcap^fl/fl^* fetal livers when compared to other genotypes **(Figures 4A-B**; **Supplemental Figure 3A**). Interestingly, surface Sca-1 expression was increased in *Srcap* knockout HSCs (**Figure 4A**). To avoid potential confounding issues secondary to Sca-1 marker dysregulation, we also classified phenotypic HSCs with Sca-1-independent methods. Using the previously characterized EPCR^32^ (**Figure 4C**; **Supplemental Figures 3B-C**) and CD86^33^ (**Supplemental Figures 3D-F**) markers, we similarly identified a similar number of phenotypic fetal liver HSCs in embryos with and without *Srcap*. When assessed at this time point, hematopoietic progenitor populations lacking in *Srcap* exhibited increased cellular proliferation as assessed by Ki-67 and 7-AAD staining when compared to corresponding cells with normal expression of *Srcap* or *Srcap* haploinsufficiency (**Supplemental Figure 3G**). This was particularly apparent in the immature CD150^+^CD48^−^ KLS compartment and may be secondary to the need to compensate for the previously demonstrated decline in more mature hematopoietic populations associated with the loss of *Srcap*.

**Figure 4.**
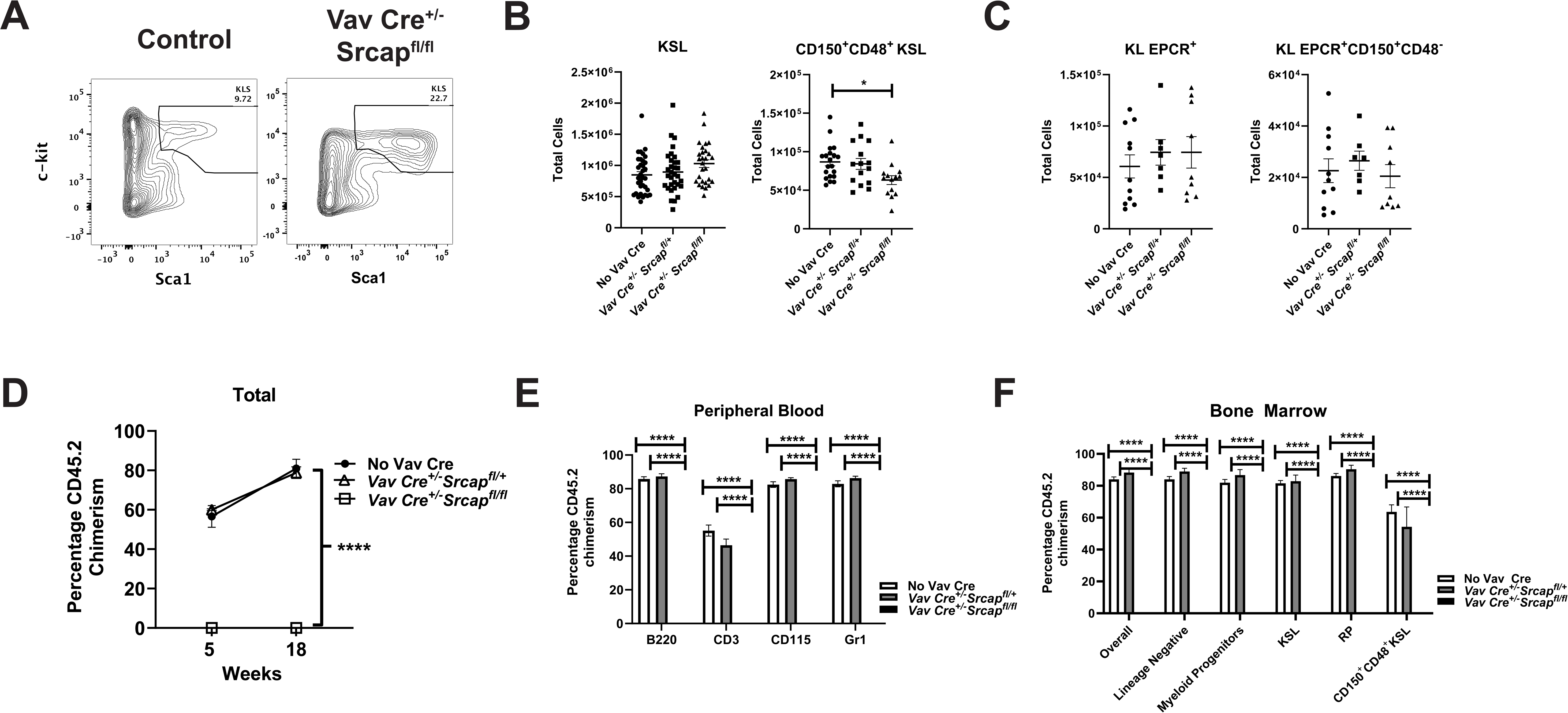
Loss of *Srcap* abrogates fetal HSC function. (A) Representative flow plots of fetal liver HSPCs using c-kit and Sca1 as markers. Cells were initially gated as CD45^+^ and lineage negative. (B) Total number of phenotypic fetal liver HSPCs using c-kit, Sca1, CD150 and CD48 as markers. (C) Total number of phenotypic fetal liver HSPCs using c-kit, EPCR, CD150 and CD48 as markers. (D) Percentage donor (Ly5.2) chimerism in the peripheral blood following competitive transplantation of 1×10^6^ Ly5.2 fetal liver cells of the indicated genotypes against 2×10^6^ wild-type Ly5.1 bone marrow cells. Vav Cre^+/−^Srcap^fl/fl^ cells were pooled from multiple embryos prior to transplantation (8 recipients in two individual experiments). Cells from embryos with full *Srcap* expression (6 donors) or haploinsufficient loss of *Srcap* (3 donors) were transplanted individually. (E) Percentage donor (Ly5.2) chimerism in the peripheral blood myeloid (Gr1^+^ and CD115^+^) and lymphoid (B220^+^ and CD3^+^) compartments 18 weeks following transplantation of fetal liver cells. (F) Percentage bone marrow donor chimerism 18 weeks following transplantation of fetal liver cells. * P ≤ 0.05. **** P ≤ 0.0001. Significance for **D** was determined with a two-way ANOVA. Significance was otherwise determined with an ordinary one-way ANOVA with multiple comparisons.

We next analyzed the functional properties of fetal liver HSPCs lacking *Srcap* by assessing their ability to repopulate following transplantation. HSPCs with normal *Srcap* expression and HSPCs with *Srcap* haploinsufficiency showed a similar ability to compete against wild-type adult bone marrow HSCs in transplantation assays. In contrast, HSPCs lacking *Srcap* failed to repopulate lethally irradiated recipients both at early and later time points (**Figure 4D**) with a failure to contribute to either the myeloid or lymphoid lineages (**Figure 4E**). Following transplantation, *Srcap* deficient HSPCs also did not reconstitute any of the recipient hematopoietic progenitor populations (**Figure 4F**). Thus, although phenotypic HSCs are present in *Srcap* deficient fetal livers, they are functionally impaired.

### Loss of *Srcap* abrogates adult HSC function

Next, we assessed whether adult HSCs recapitulated the phenotypes of their fetal liver counterparts. *Mx1-Cre^+/−^Srcap^fl/fl^*mice treated with poly(I:C) began exhibiting signs of illness and died within two-three weeks of drug administration (**Figure 5A**). *Mx1-Cre^+/−^Srcap^fl/fl^* mice sacrificed at the first sign of illness displayed a decline in peripheral blood counts (**Figure 5B**) and a progressive loss of bone marrow c-kit^+^ cells (**Figure 5C**), suggesting that *Srcap* plays an essential role in maintaining adult hematopoiesis. To determine if this role is cell-intrinsic in nature, we non-competitively transplanted 5×10^6^ bone marrow cells with inducible loss of *Srcap* (*Mx1-Cre^+/−^Srcap^fl/fl^*; *Cre^+^*) or control (*Srcap^fl/fl^*) bone marrow cells into lethally irradiated recipients followed by poly(I:C) treatment 5-6 weeks post-transplantation. As expected, control bone marrow cells efficiently repopulated the recipient hematopoietic compartment. However, *Cre^+^*bone marrow cells failed to reconstitute hematopoietic populations in recipient mice (**Figure 5D**). This included both the myeloid and lymphoid compartments (**Supplemental Figure 4A**) and the bone marrow hematopoietic progenitor populations (**Figure 5E**). Of note, we observed an initial decline in *Cre^+^* repopulation even before the administration of poly(I:C), potentially due to leakiness of the Mx1-Cre transgene caused by post-transplantation inflammation. These data show that hematopoietic-intrinsic *Srcap* is essential for the proper function of adult HSCs.

**Figure 5.**
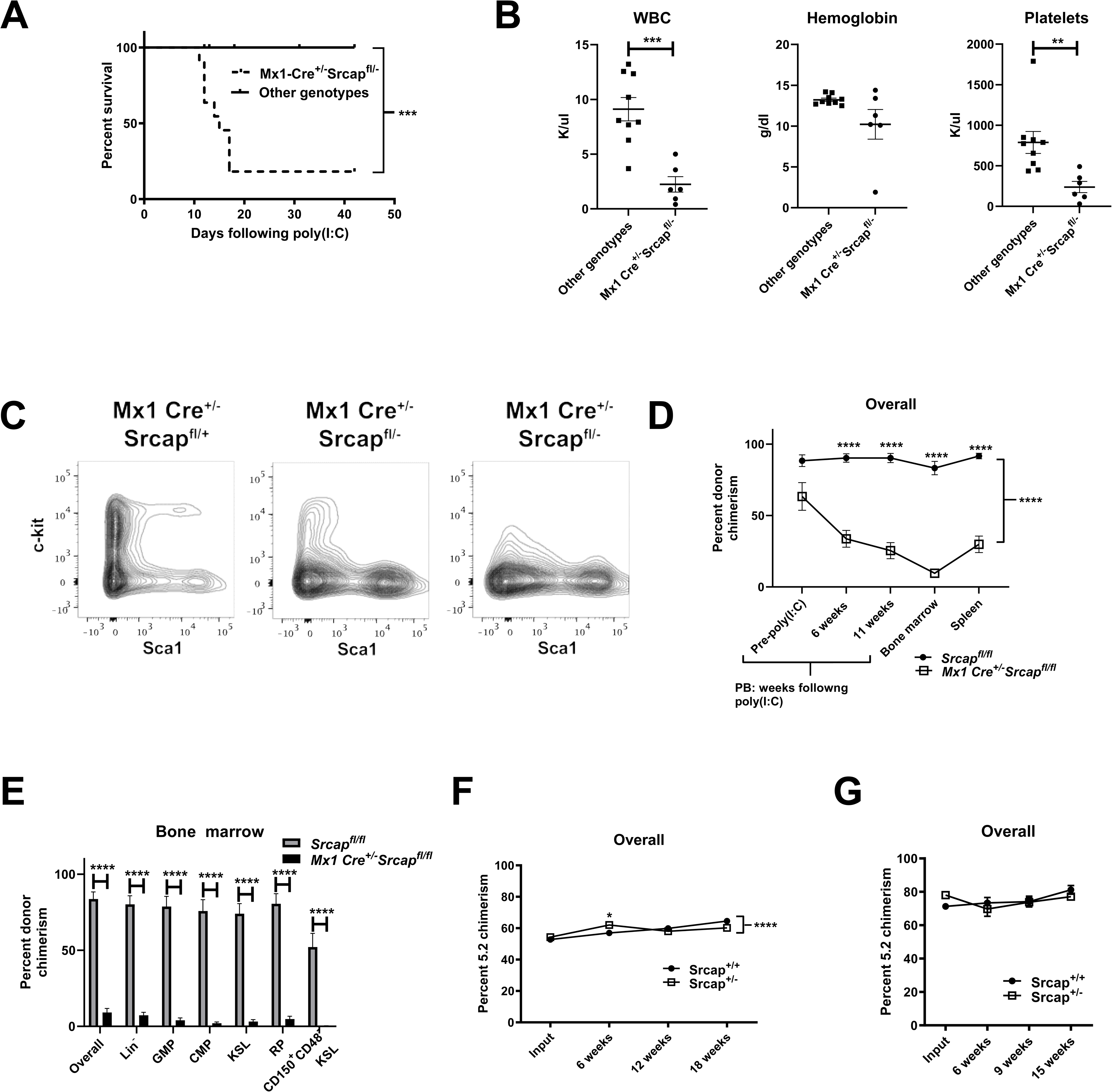
Loss of *Srcap* abrogates adult HSC function. (A) Survival of mice following poly(I:C) administration to induce loss of *Srcap*. Survival was measured from the date of the last dose of poly(I:C), and significance assessed with a Mantel-Cox test. (B) Hematopoietic parameters following poly(I:C) administration. Mx1 Cre^+/−^ Scrap^fl/-^ mice were analyzed upon exhibiting signs of distress with control mice analyzed at similar time points. Significance was assessed with an unpaired t-test. (C) Representative bone marrow flow plots following poly(I:C) administration. Cells were initially gated as lineage negative. (D) Percentage donor (Ly5.2) chimerism in the indicated tissues following non-competitive transplantation of 5×10^6^ Mx1 Cre^+/−^ Srcap^fl/fl^ or Srcap^fl/fl^ bone marrow cells (8 recipients per group in two independent experiments) and poly(I:C) administration. Significance was assessed with a two-way ANOVA. (E) Percentage bone marrow donor chimerism 11-12 weeks following poly(I:C) administration with significance assessed using an unpaired t-test. (F) Percentage peripheral blood donor (Ly5.2) chimerism following competitive transplantation of 2.5×10^6^ Srcap^+/−^ or Srcap^+/+^ bone marrow cells against an equal number of competitor cells (14-18 recipients per group in 2 independent experiments). Significance was assessed with a two-way ANOVA. (G) Percentage peripheral blood donor (Ly5.2) chimerism after re-transplantation of Srcap^+/−^ or Srcap^+/+^ bone marrow cells (5 recipients per group). The input chimerism is the chimerism of the bone marrow KLS cells at the time of re-transplantation. Significance was assessed with a two-way ANOVA. ** P ≤ 0.01. *** P ≤ 0.001. **** P ≤ 0.0001.

In clonal hematopoiesis and in myeloid malignancies, *SRCAP* mutations are heterozygous in nature. However, fetal liver hematopoiesis is normal in *Vav1-iCre^+/−^Srcap^fl/+^*mice, with no significant defects either at steady state or after transplantation (see **Figures 2-4**). We next determined whether the same was true for adult HSCs by competing bone marrow cells from mice with constitutive germline *Srcap* haploinsufficiency (*Srcap^+/−^* genotype) against an equal number of age and sex-matched wild-type competitors in transplantation assays. Like fetal liver hematopoietic cells, adult bone marrow cells with heterozygous loss of *Srcap* maintained an overall normal repopulating capacity both in primary (**Figure 5F**) and secondary (**Figure 5G**) transplantation experiments. Interestingly, *Srcap^+/−^* bone marrow cells showed a similar ability to repopulate the myeloid compartment as wild-type control cells after both primary and secondary transplantation. In contrast, a modest advantage in repopulating the lymphoid compartment after primary transplantation (**Supplementary** Figure 4B) was observed, which does not persist after secondary transplantation (**Supplementary** Figure 4C). This is similar to the phenotype observed upon transplanting HSCs with heterozygous expression of an *Srcap* truncation mutant allele.^34^

Collectively, these data show that adult and fetal liver hematopoiesis is maintained in *Srcap* haploinsufficient mice, with no evidence that *Srcap* haploinsufficiency confers a fitness advantage to HSCs.

### Cellular impacts of *Srcap* loss

Finally, we investigated the transcriptomic consequences of *Srcap* loss. We focused on fetal liver stage 1 erythroid progenitors, which is the stage prior to where we observed a significant loss in the erythroid population. We used RNA-Seq to compare cells with normal *Srcap* expression, heterozygous loss of *Srcap*, and complete loss of *Srcap*. Consistent with the phenotypic consequences of *Srcap* deficiency, cells with normal *Srcap* expression or heterozygous loss of *Srcap* were similar transcriptionally, while both were distinct from cells having complete *Srcap* loss. (**Figure 6A**). The top upregulated gene expression pathway in *Srcap* deficient erythroid cells was the p53 pathway **(Figures 6B-C**; **Supplemental Table 1**). Other top hits included inflammatory signaling and apoptosis pathways. Interestingly, *Srcap* deficient fetal liver hematopoiesis was associated with increased CD44 erythroid progenitor (**Figure 2C**) and Sca1 HSPC (**Figure 4A**) surface expression, consistent with the reported increase in these two surface markers following exposure to inflammatory signaling.^35,36^

**Figure 6.**
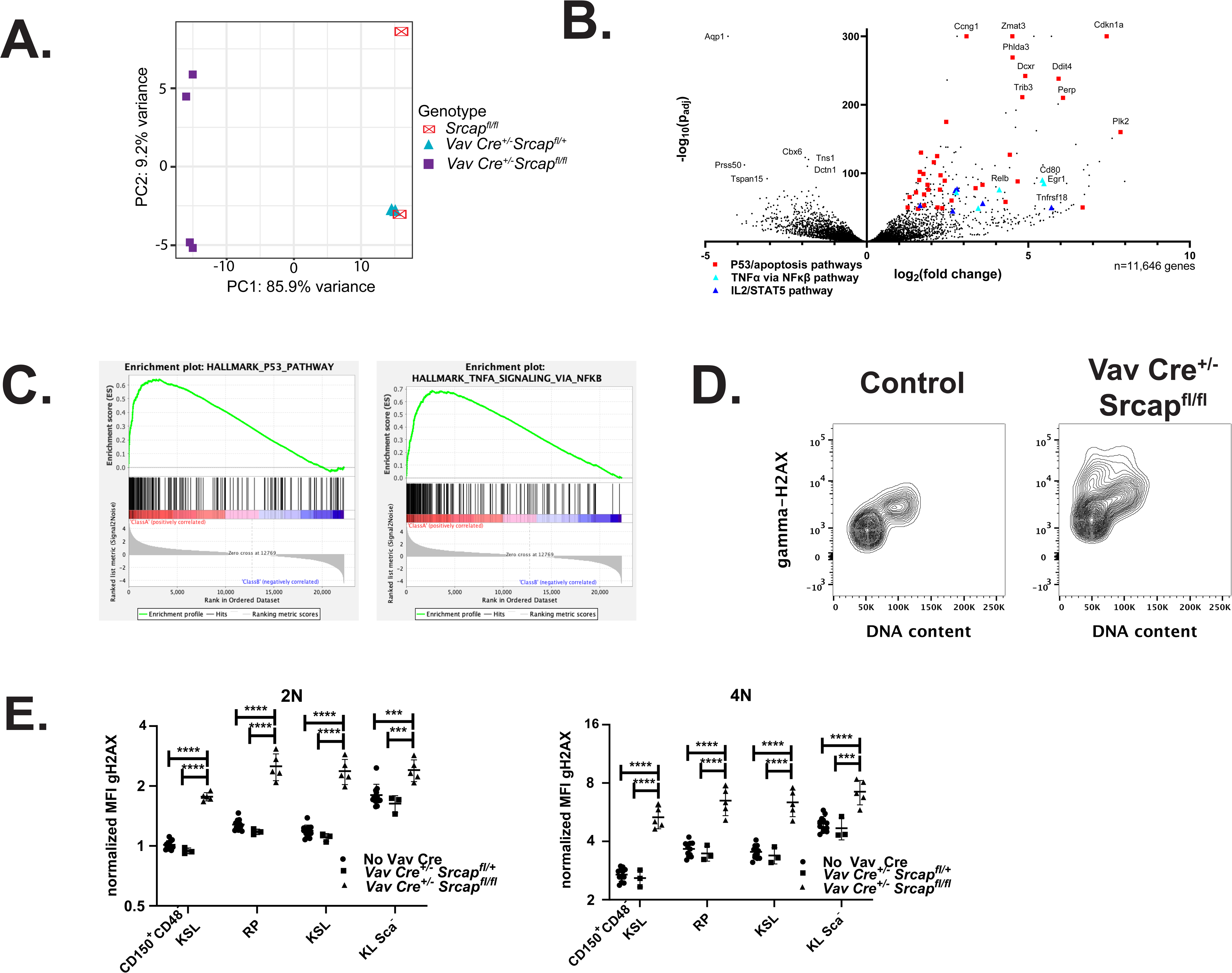
Cellular impacts of *Srcap* loss. (A) Principal component analysis of RNA-Seq data from stage 1 fetal liver erythroid progenitors from *Vav Cre^+/−^*Srcap^fl/fl^ embryos (n=4), *Vav Cre^+/−^* Srcap^fl/+^ embryos (n=2) and Srcap^fl/fl^ embryos (n=2). (B) Volcano plot of differentially expressed genes in RNA-Seq data of stage 1 fetal liver erythroid progenitor cells with complete loss of *Srcap* (n=4) versus cells with either full or heterozygous loss of *Srcap* expression (n=4). Genes with mean DeSeq2 values > 50 are shown. Genes are highlighted if they are in the P53, apoptosis, TNFα via NFκβ, or IL2/STAT5 GSEA Hallmark pathways and among the top 250 most significantly dysregulated genes. Genes highlighted in the latter two pathways were not also present in the first two. (C) P53 pathway and TNFα via NFκβ GSEA enrichment plots from the analysis in **B**. These had the two highest normalized enrichment scores among the GSEA Hallmark pathways. (D) Representative flow plots showing γ-H2AX levels in CD150^+^CD48^−^ KSL cells. DNA content was assessed with 7-AAD staining. (E) γ-H2AX MFI in the fetal liver hematopoietic progenitor compartment. The MFIs were normalized to the average MFI of control CD150^+^CD48^−^ KLS cells having 2N DNA content with either full or heterozygous loss of *Srcap* expression. Significance was determined with an ordinary one-way ANOVA with multiple comparisons. *** P ≤ 0.001. **** P ≤ 0.0001.

SRCAP has previously been proposed to play a direct role in DNA double-strand break repair,^37^ and the presence of a commonly identified *Srcap* truncation mutation was recently demonstrated to alter the expression of multiple genes involved in the repair of DNA damage.^34^ Thus, we assessed if the transcriptional upregulation of p53 target genes associated with *Srcap* loss was accompanied by evidence of increased DNA damage, focusing on the fetal liver HSPC compartment due to the robust DNA damage machinery maintained in fetal liver HSCs.^38^ In control embryos, γ-H2AX median fluorescent intensity (MFI) increased as hematopoietic progenitors matured from HSCs to more differentiated progenitors and was higher in cells in the G2M phase of the cell cycle versus cells with 2N DNA content (**Figure 6D-E**), consistent with increased γ-H2AX levels in actively replicating cells.^39^ Compared to controls, *Vav1-iCre^+/−^ Srcap^fl/fl^* hematopoietic progenitors had an increased γ-H2AX MFI in all assessed stages of hematopoietic progenitor maturation and in cells with either 2N or 4N DNA content, with the increase particularly prominent in the immature HSC compartment. Collectively, these data suggest that the loss of *Srcap* is associated with evidence of increased DNA damage and concurrent activation of the p53 pathway.

## Discussion

In this study, we generated a conditional *Srcap* loss of function murine model to determine the functional consequences of hematopoietic-specific loss of this gene. We discovered that hematopoietic-specific *Srcap* loss is embryonic lethal. Certain hematopoietic populations were particularly sensitive to *Srcap* loss with its ablation resulting in the early decline of fetal liver erythroid progenitor and monocytic populations while phenotypic granulocytic and HSPC populations were initially preserved. However, fetal liver HSPCs lacking *Srcap* failed to repopulate in bone marrow transplantations assays, and adult HSPCs with induced loss of *Srcap* failed to maintain hematopoiesis, indicating that this gene is essential for both hematopoietic maturation and HSPC function.

Indeed, AML-associated mutations in *SRCAP* appear to be exclusively heterozygous in nature,^23^ suggesting that, in hematopoietic populations harboring such mutations, the function of SRCAP is perturbed, but not completely eliminated. Supporting this, previous studies have demonstrated that alterations in Srcap activity, whether through loss of the Znhit1 subunit of the Srcap complex^15,19^ or via expression of a truncated Srcap mutant,^34^ can impact the normal functioning of HSPCs. In our murine model of *Srcap* loss, heterozygous loss of this gene does not adversely impact HSPC repopulating ability, indicating that hematopoietic progenitors that maintain at least one wild-type *SRCAP* allele remain capable of sustaining hematopoiesis. Interestingly, similar to a recently published murine model with expression of a commonly observed *Srcap* truncation mutant,^34^ hematopoietic bone marrow cells with heterozygous loss of *Srcap* displayed a lymphoid bias after transplantation. How the different *SRCAP* frameshift and nonsense mutations observed in human hematopoiesis impact SRCAP activity (e.g. the degree to which they abrogate wild-type function through loss of the wild-type allele or partial dominant negative activity or provide a novel gain of function) remains to be determined.

To investigate the cellular consequences of SRCAP deficiency in hematopoietic populations, we performed RNA-Sequencing on *Vav1-iCre^+/−^Srcap^fl/fl^* fetal liver stage 1 erythroid progenitors. With complete loss of *Srcap*, this population was associated with the significant upregulation of p53, apoptotic, and inflammatory pathways. Interestingly, shRNA-induced knockdown of SRCAP in a non-hematopoietic cell line has been similarly associated with a more modest upregulation of inflammatory pathways and the p53 pathway.^40^ Upregulation of such pathways may be common downstream results of *SRCAP* loss in hematopoietic populations, providing a potential explanation as to why some degree of *SRCAP* expression is required to maintain hematopoiesis. It is also possible that the significant upregulation of p53 and inflammatory pathways associated with complete *Srcap* loss is masking milder impacts loss of this gene has on other cellular pathways. Indeed, previous publications have also identified both SRCAP and H2AZ as playing a critical role in the cellular response to DNA damage.^34,37,41^ The degree to which milder perturbations of SRCAP impact such pathways in human clonally expanded and/or leukemic populations harboring *SRCAP* mutations, the dependence of these impacts on mutation-induced alterations to SRCAP’s function in the chromatin deposition of H2AZ, and the mechanisms through which this may promote clonal expansion and leukemic transformation all remain open areas of investigation.

## Acknowledgements

This work was supported by National Institutes of Health, National Cancer Institute grant K08 CA197369 (to T.N. Wong) and by the Leukemia Research Foundation. At Washington University School of Medicine, technical support was provided by the Transgenic, Knockout and Micro-Injection and High-Speed Cell Sorting Cores. At the University of Michigan, technical support was provided by the Flow Cytometry, Advanced Genomics, and Bioinformatics Cores.

## Authorship

Contributions: T.N.W. and D.C.L. initiated the project, designed the research, and wrote the paper with input from other authors. T.N.W., A.M., E.R.F., D.K., and A.C. performed the research and analyzed the data.

## Figure Legends

**Supplemental Figure 1. Loss of *Srcap* causes aberrant hematopoietic maturation.** (A) Representative H&E stains of fetal livers with the indicated genotypes. (B) Percentage of fetal liver cells that were Ter119^+^. (C) Percentage of fetal liver erythroid progenitors at the indicated stage of maturation based on forward scatter, Ter119 staining, and CD44 staining.

**Supplemental Figure 2. Loss of *Srcap* causes aberrant fetal myeloid maturation.** (A) Percentage of fetal liver cells classified as phenotypic CD11b^+^CD115^−^ granulocytes or CD11b^+^CD115^+^ monocytes as described in **Fig. 3A**. (B) Percentage of fetal liver cells classified as Ly6C^Int^CD11b^+^ granulocytes or Ly6C^High^CD11b^Int^ monocytes as described in **Fig. 3A**. (C) Representative flow plots identifying fetal liver myeloid populations using the markers CD11b, Ly6G, F4/80, and CD115. (D) Total number of fetal liver cells classified as phenotypic CD11b^+^Ly6G^+^ granulocytes or F4/80^+^CD115^Int^ macrophages as described in **C**. (E) Percentage of fetal liver cells classified as phenotypic CD11b^+^Ly6G^+^ granulocytes or F4/80^+^CD115^Int^ macrophages as described in **C**. (F) Percentage of phenotypic CMPs, GMPs, and MEPs among fetal liver cells. ** P ≤ 0.01. *** P ≤ 0.001. **** P ≤ 0.0001. Significance was determined with an ordinary one-way ANOVA with multiple comparisons.

**Supplemental Figure 3. HSCs with loss of *Srcap* are initially maintained in the fetal liver.** (A) Percentage of fetal liver cells classified as phenotypic HSPCs using c-kit, Sca1, CD150 and CD48. (B) Representative flow plots of fetal liver HSPCs using c-kit and EPCR as markers. Cells were initially gated as CD45^+^ and lineage negative. (C) Percentage of fetal liver cells classified as phenotypic HSPCs using c-kit, EPCR, CD150 and CD48. (D) Representative flow plots of fetal liver HSPCs using c-kit and CD86 as markers. Cells were initially gated as CD45^+^ and lineage negative. (E) Total number of phenotypic fetal liver HSPCs using c-kit, CD86, CD150 and CD48. (F) Percentage of fetal liver cells classified as HSPCs using c-kit, CD86, CD150 and CD48. (G) Cell-cycle distribution of fetal liver cells. 2N, Ki-67 low cells were classified as in G_0_, 2N, Ki-67 high cells were classified as in G_1_, and 4N cells were classified as in G_2_-M/S. * P ≤ 0.05. ** P ≤ 0.01. *** P ≤ 0.001. **** P ≤ 0.0001. Significance was determined with an ordinary one-way ANOVA with multiple comparisons.

**Supplemental Figure 4. Loss of *Srcap* abrogates HSC function.** (A) Percentage peripheral blood donor (Ly5.2) chimerism in the myeloid (Gr1^+^ and CD115^+^) and lymphoid (B220^+^) lineages following non-competitive transplantation of 5×10^6^ Mx1 Cre^+/−^Srcap^fl/fl^ or Srcap^fl/fl^ bone marrow cells. (B) Percentage peripheral blood donor (Ly5.2) chimerism in the myeloid (Gr1^+^ and CD115^+^) and lymphoid (B220^+^) lineages following competitive transplantation of 2.5×10^6^ Srcap^+/−^ or Srcap^+/+^ bone marrow cells against an equal number of competitor cells. (C) Percentage peripheral blood donor (Ly5.2) chimerism in the myeloid (Gr1^+^ and CD115^+^) and lymphoid (B220^+^) lineages after re-transplantation of Srcap^+/−^ or Srcap^+/+^ bone marrow cells. Significance was assessed with a two-way ANOVA. * P ≤ 0.05. *** P ≤ 0.001. **** P ≤ 0.0001.

**Supplemental Table 1.** Upregulated gene pathways in stage 1 fetal liver erythroid progenitors with loss of *Srcap*.

