## Supplementary figures and images for "The Clonal Hematopoiesis-associated Gene *Srcap* Plays an Essential Role in Hematopoiesis"

### Supplemental Figures

#### SUPPLEMENTAL FIGURE 1

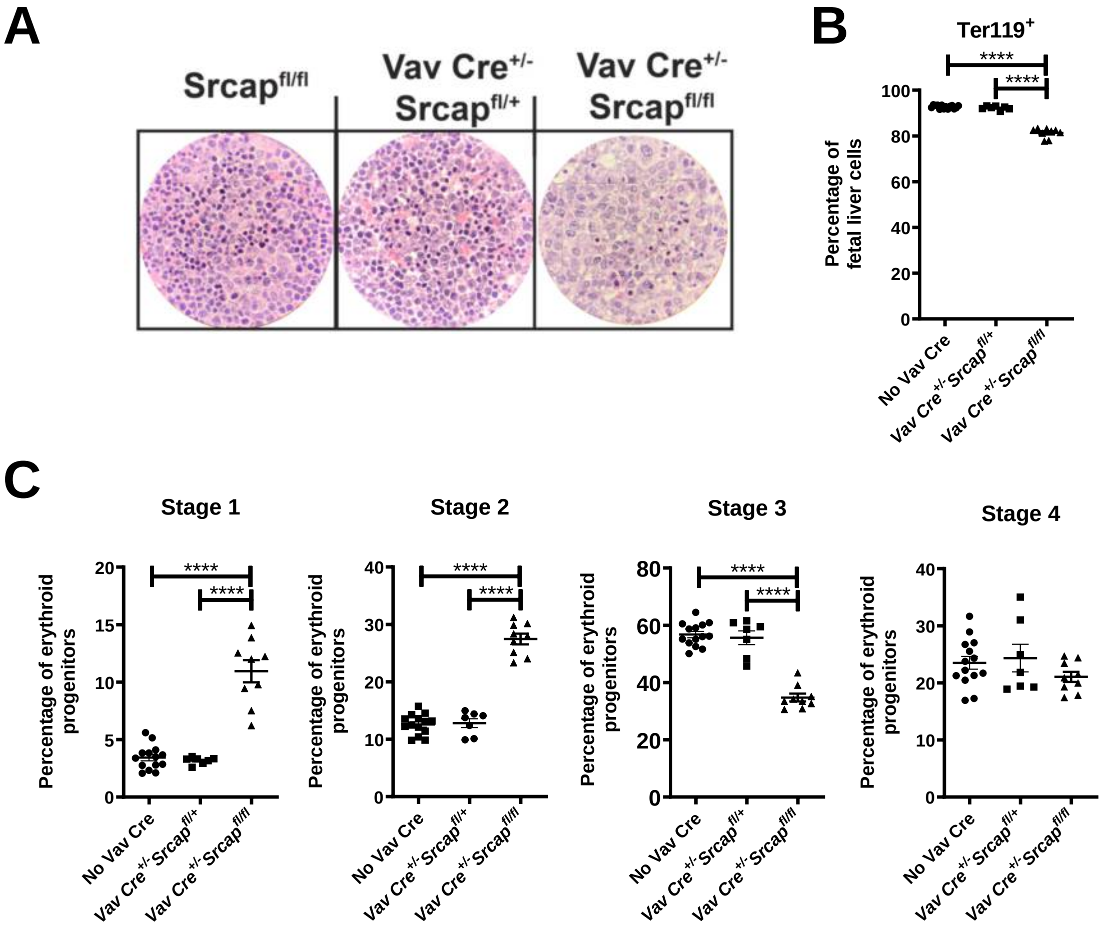

#### Supplemental Figure 2

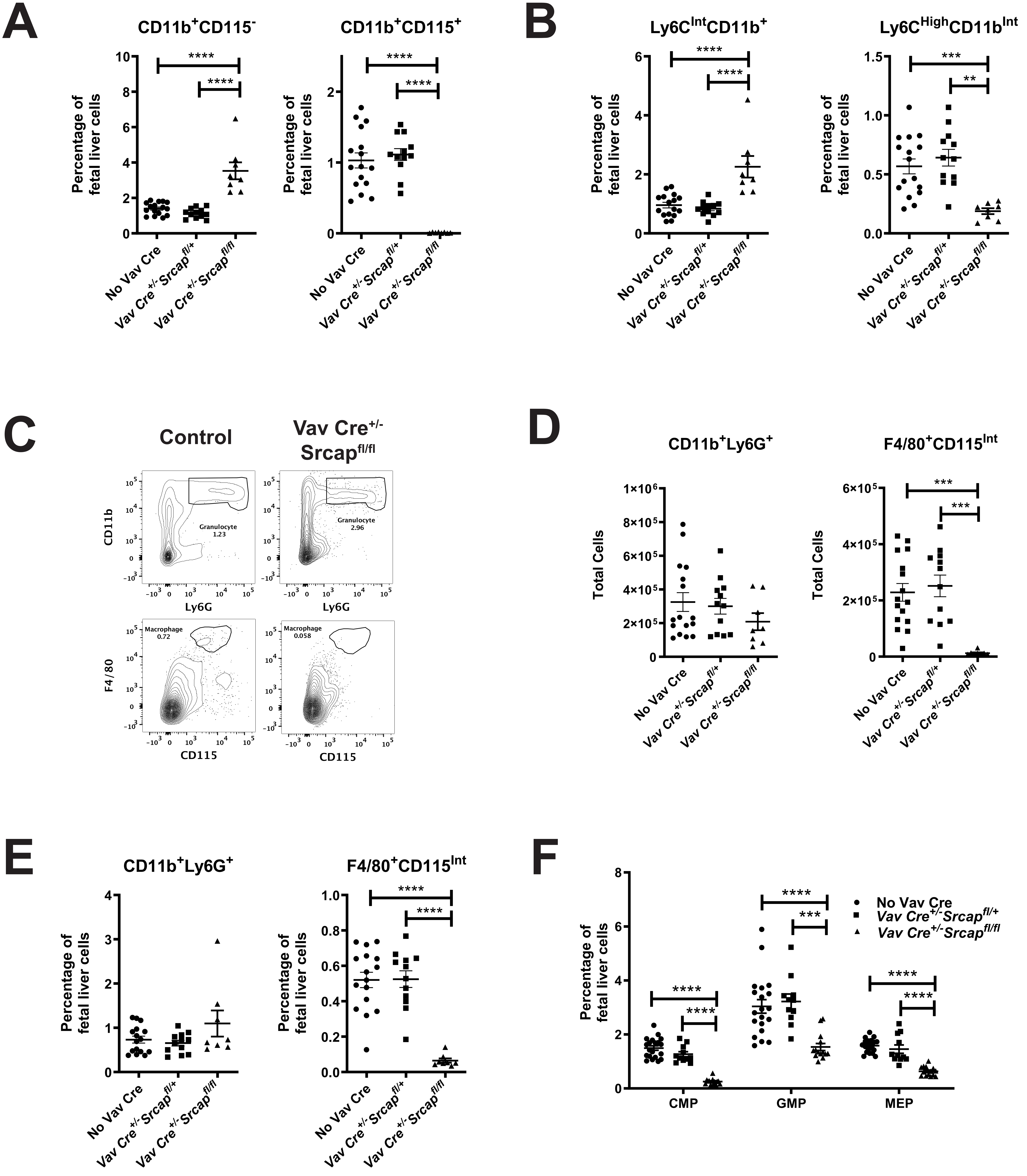

#### Supplemental Figure 3

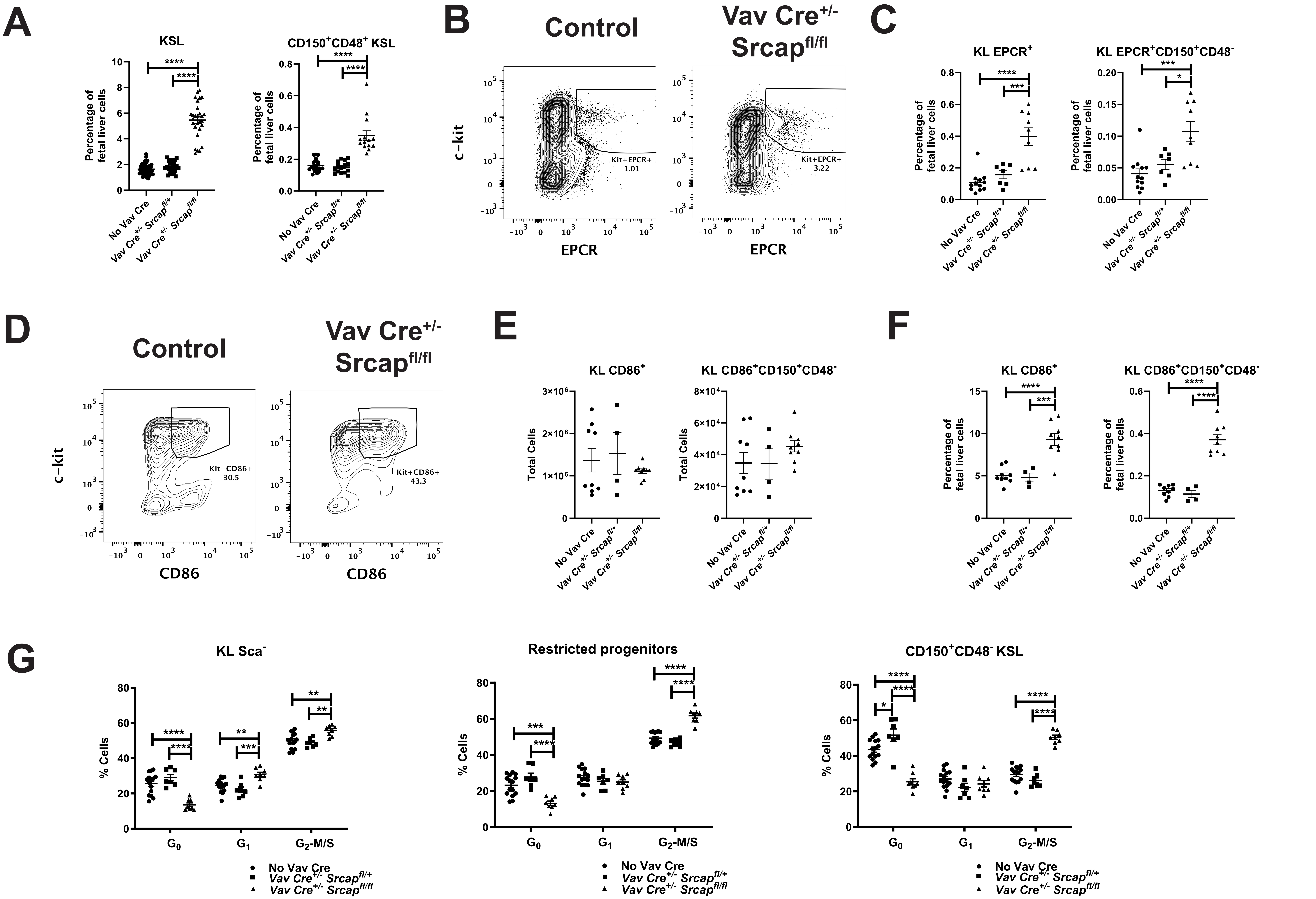

#### SUPPLEMENTAL FIGURE 4

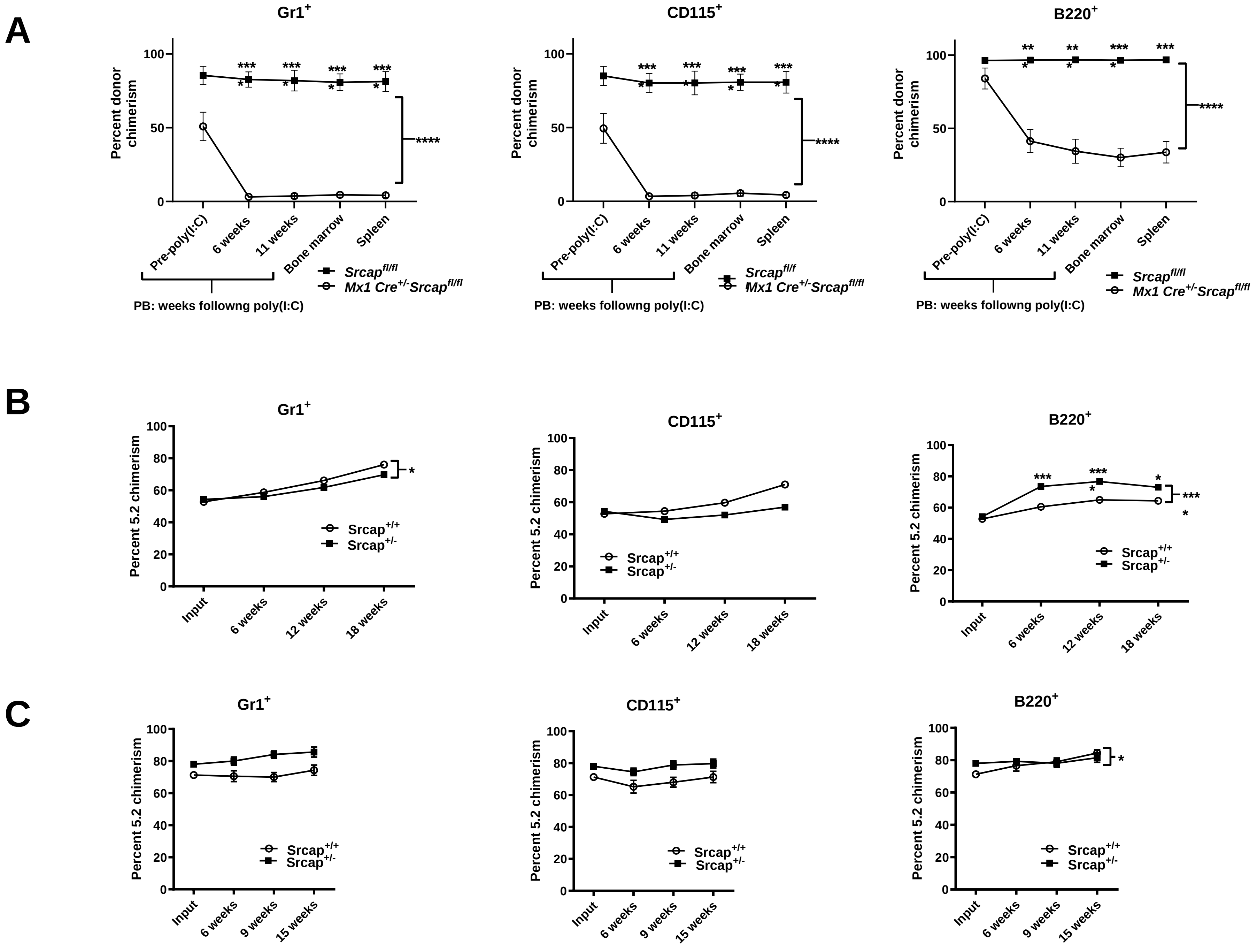
